# Combining protein-based transcriptome assembly, and efficient MinION long read sequencing for targeted transcript sequencing in orphan species. Validation on herbicide targets and low copy number genes in Gymnosperms, Juncaceae and Pteridophyta

**DOI:** 10.1101/2020.10.24.353441

**Authors:** Dyfed Lloyd Evans

## Abstract

Orphan species that are evolutionarily distant from their closest sequenced/assembled neighbour provide a significant challenge in terms of gene or transcript assembly for functional analysis. This is because 30% sequence divergence from the closest available reference sequence means that, even with a complete genome or transcriptome sequence, mapping-based or reference-based approaches to gene assembly and gene identification break down.

A new approach is required for reference-guided gene and transcript assembly in such orphan species, or species that are evolutionarily very divergent from their closest relatives. When annotating genes, the protein sequence is often preferred as it diverges less than the DNA/RNA sequence and it is often simpler to find meaningful homology at the protein level. This greater conservation of protein sequence across evolutionary time also makes proteins a prime candidate for use as the basis for sequence assembly. A protein-based pipeline was developed for transcript assembly between distantly related species. This was tested on three evolutionarily divergent species with little sequence information available for them and for which the closest genome representatives were at least 40 million years divergent as well as one species (*Azolla filiculoides*) for which a genome assembly is available. All the species have the potential to be weeds and herbicide targets were chosen as functional genes, whilst low copy number genes were chosen for evolutionary studies. Transcriptomic sequences were assembled using a bait and assemble strategy and final assemblies were verified by direct sequencing.

## Introduction

Genome analysis in orphan species (and crops) can often be very challenging. Considerably less funding is available and research groups are often smaller. Orphan species are also typically quite distantly related to species for which extensive genome and transcriptome datasets are available. The challenge for orphan species is the development of cost-effective tools and techniques that can be applied on commodity hardware. This includes both bioinformatics tools and sequencing technologies.

Though great advances are being made in *de novo* assembly of transcriptomic datasets, assembly of such data from short read datasets is still a hard problem (Góngora-Castillo et al. 2013). Even more so if only a limited depth of sequencing is available, as this tends to produce large amounts of fragmented transcripts (Bens et al. 2016). Though long read technologies such as PacBio Sequel and Oxford Nanopore Technologies MinION allow for the capture of full-length transcripts these technologies remain comparatively expensive and are prone to high error rates (Pollard et al. 2018). All these technologies, in *de novo* assemblies also reveal large amounts of sequence contamination from bacterial and fungal metabiome associations as well as human contamination from sample handling (Merchant et al. 2014). As a result, whole-scale assembly of these sequences can waste effort in terms of contaminant identification and removal.

Optimal transcriptome and genome assemblies are still attained though reference- guided assembly. However, at the DNA level this is not viable beyond about 30% sequence variation (Ungaro et al. 2017). In plants this typically equates to 15–20 million years divergence. Beyond this point it is not possible to perform a reference-guided assembly (Parchman et al. 2010) or to use bait-and-assemble methodologies (Lloyd Evans and Joshi, 2016a; Lloyd Evans and Joshi 2017) to assemble a sequence based on a reference.

In contrast, genome annotation is often performed at the protein level — as proteins typically diverge less than DNA over the same periods of time (Huerta-Cepas et al. 2015). Effectively, proteins retain more sequence relations (and thus functional relations) despite the divergence of their parent species. The recent development of Diamond (Buchfink et al. 2015) a fast DNA to protein mapper that works with short reads means that the development of a pipeline that uses the protein of one species to assemble transcripts from a much more divergent species is possible.

Such a pipeline was developed employing Diamond, initially to assemble the six major herbicide target genes in four species, but which was then extended to assemble 14 evolutionarily informative low copy number genes across 16 species. Assembled transcript sequences were confirmed by PCR amplification and MinION sequencing.

*Pteridium aquilinum* (known as bracken or bracken fern) has a global distribution, and counts amongst the top five most numerous vascular plants worldwide. It is a perennial member of the pteridophyte (fern) family and includes numerous sub-species and varieties. It is most common in semi-shaded well-drained open woodlands or woodland margins, though it also thrives on peat-based moorland. It predominantly spreads through dense rhizomatous networks but can re-establish through spores.

Bracken can be a major weed in grazing grassland, particularly in reclaimed land. In addition, due to the presence of norsesquiterpene glycosides (ptaquiloside, ptesculentoside, and caudatoside) which cause haemorrhagic disease, particularly in ruminants (Vetter 2009). Acute brackenism occurs when animals ingest high doses over relatively short durations of weeks or months and is characterized by bleeding. Of the various toxic chemicals found in bracken, the one of most concern is ptaquiloside, which can cause neoplasias and is a known carcinogen. In sheep, ptaquiloside poisoning can cause bright blindness (Hirono et al. 1993). In monogastric animals, such as equines, acute poisoning typically results in bracken staggers, resulting from thiamine deficiency (Taylor 1980).

Moreover, epidemiological studies suggest that ptaquiloside should be of concern to humans (Pamukcu et al. 1978). Consumption of milk from cattle with access to bracken increases the risk of human oesophageal and gastric cancer. Japanese and Welsh research have also shown an association between consumption of bracken crozier and oesophageal cancer (Galpin et al. 1990). Additionally, ptaquiloside has been found as an environmental contaminant in soil and water, and air-borne spores may also present a risk of human exposure (Rasmussen et al. 2005; Clauson-Kaas et al. 2014).

Bracken growth can be retarded by close grazing or trampling in alternative grazing pasture systems. Fern density can be reduced by regular cutting of the mature plant or, if the land is suitable, by deep ploughing. Herbicide treatment using glyphosate and asulam (a 7,8- dihydropterate synthase inhibitor and a cell division inhibitor) can be an effective method of control, especially if combined with cutting before treatment (Pakeman et al. 1997).

*Equisetum arvense*, the field or common horsetail, is a primitive spore-bearing plant in the Equisetaceae family. It is a common weed in the UK and elsewhere, particularly in fruit (and other perennial corps) in wheat and in nurseries (Lipecki 2006). It readily accumulates heavy metals and is toxic to sheep, cattle and horses even when dried as hay. The causal agent of such toxicity is believed to be thiaminase — either acting alone or in conjunction with one or more additional toxins (Hill and Foland, 1986). It is botanically of interest as it contains natural fungicides, which can be used as an extract of the plant (Garcia et al. 2011). It is a rhizomatous perennial and is considered weedy. It has deep freely-branching rhizomes with round tubers. It matures and spreads quickly, forming dense, long-lived mats, tolerates flooding to dampish-dry soil, warm to very cold, wind, and deep burial but is intolerant of dry soils. The plant can re-grow from only a small section of the rhizome or tuber and as these can extend to 1.8m underground herbicide control can be difficult (Marshall 1986). Ploughing or rotovating affords no control and will even spread the plant more effectively (this is why it’s particularly troubling in nurseries).

For control, small patches of the plant can be dug out and Primisulfuron (Beacon) gives reasonable burning action on horsetail in grain crops. Repeated applications of MCPA (a phenoxy herbicide inhibiting photosynthesis) reportedly reduces horsetail infestations and would be safe to pasture grasses (Soltani et al. 2015). The Weed Control Manual 2000 (Curran, et al., 2000) lists only two herbicides for field horsetail control in non-cropland, ornamentals/woody plantings, small fruits and deciduous tree fruits: diclobenil (Casoron) for all of the above sites/crops; and clorsulfuron (Telar) or sulfometuron (Oust) for non-crop areas. In the UK, glufosinate-ammonium is typically used to eradicate horsetail, particularly in horse pastureland (Lipecki 2006). This is a non-selective contact herbicide that inhibits glutamine synthetase.

*Azolla filliculoides* is a floating aquatic fern native to warm temperate and tropical regions of the Americas. It can fix nitrogen from the air, and as a result has been introduced into Western Europe, southern Africa, tropical Asia, Australia and New Zealand where, in certain circumstances, it has become an invasive weed. Azolla filliculoides was first recorded in the UK in 1883 and is considered invasive in southern England and coastal Wales (NNSS UK 2015). Due to its invasive properties it was decided to analyse the plant for herbicide target genes for potential future use. *Azolla filliculoides* is one of two fern genomes to have been sequenced (Li et al. 2018) and could provide a reference against which the sequences derived from this project could be independently verified.

*Juncus effusus* (common rush or soft rush) is a perennial flowering plant in the Juncaceae family. It is a member of the Poaceae that is considered native in Europe, Asia, Africa, North America, and South America. It typically grows in wetlands, riparian areas, ditches, and marshes. In the UK it is typically found in rush pastures and fen-meadow environments and is typically associated with peat grasslands (Rodwell 1991). It is generally unpalatable to grazing animals and is considered a weed of pasture-land. In terms of herbicides, it can be controlled by 2,4-D and imazaquines (typically Imazamox in the UK). It is considered a major weed of establishing coniferous trees in cut-away peatland (McCrory and Renou 2003) *Juncus effusus* is a member of the Juncaecae, within the Poales, and is ancestral to true grasses, but 20 million years divergent from them (Jones et al. 2007).

Equisetales have recently been placed as a member of the pteridophytes and sister to the remaining fern families (Shen et al. 2017). *Pteridium aquilinum* is a member of the Dennstaeditioids and is sister to the core Eupolyploids. Genomes from two members of the early leptosporangiates, *Azolla filiculoides*. and *Salvinia cucullata* have been sequenced, but bracken is 145 million years divergent from these species (Lie et al. 2018; Shen et al. 2017). The closest assembled genome to *Juncus effusus* is that of *Ananas comosus*, which is 120 million years divergent (Bouchenak-Khelladi et al. 2014).

*Juncus effusus*, *Pteridium aquilinum* and *Eqiusetum arvense* are therefore well outside the window where standard gene/transcript assembly by mapping would work. A new protein-based gene/transcript assembly pipeline was developed. This was employed to assemble the seven major known herbicide target genes in all species. These being: Acetyl CoA Carboxylase (ACCase: the target of aryloxyphenoxypropionate, cyclohexanedione and phenylpyrazolin herbicides) (Kukorelli et al. 2013); acetolactate synthase (ALS: the target of sulfonylureas (SUs), imidazolinones (IMIs), triazolopyrimidines (TPs), pyrimidinyl oxybenzoates (POBs), and sulfonylamino carbonyl triazolinones (SCTs)) (Zhou et al. 2007); 5-enolpyruvylshikimate-3-phosphate synthetase (EPSPS: the target of glyphosate) (SchoLnbrunn et al. 2001); Glutamine synthase 2 (GS2; the target of Glufosinate) (Berlicki 2008); 4-Hydroxyphenylpyruvate dioxygenase (HPPD: the target of pyrazolynate, pyrazoxyfen, benzofenap, sulcotrione and mesotrione) (Kraehmer et al. 2014); phytoene desaturase (PDS: the target of bleaching herbicides) (Sandmann et al. 1991) and chloroplastic Protoporphyrinogen oxidase (PPO: the target of Diphenylether, N-phenylphthalimide, Oxadiazole, Pyrimidindione, Thiadiazole and Triazolinone class herbicides) (Hao et al. 2011). As GS2 and PPO have closely-related orthologues, these were also assembled and sequenced. In addition, thirteen low copy number genes useful in phylogenetic analyses were also assembled and sequenced. Assemblies were confirmed by Oxford Nanopore Technologies MinION sequencing of concatenated PCR products corresponding to the gene assemblies using samples collected in the UK.

In grasses, low copy number genes employed for phylogenetic analyses are typically flowering genes (Lloyd Evans et al. 2019). However, horsetails and ferns have different reproductive mechanisms from both gymnosperms and angiosperms (Hasebe 1999). Previously, Zhang et al. (2012) and Zeng et al. (2014) have demonstrated the use of low copy number transcripts for deep angiosperm phylogeny. They revealed the divergence of eudicots, monocots, magnoliids, Chloranthaceae and Ceratophyllaceae. However, gymnosperms were only used as an outgroup and polypodiopsida were not included. In this paper, the utility of a subset of these genes, assembled based on protein mapping, for resolving the position of Gymnospermae and Polypodiopsida within the broader land plant phylogeny. As well as the three target species of this study, the low copy number genes from: *Ginko biloba*, an early- diverging gymnosperm; *Taxus baccata* and *Pinus sylvestris* were assembled and sequenced to fill the gaps in the early portions of the low copy number phylogeny. In addition, the low copy number transcripts of *Lygodium japonicum*, *Ophioglossum vulgatum*, *Gnetum montanum*, *Acorus calamus*, *Platanus occidentlais*, *Berberis thungbergii* and *Lilium lancifolium* were assembled from public RNA-seq data.

The data presented represent the first ever large-scale study of herbicide target genes in orphan weeds, employing a novel methodology for transcript assembly from NGS data using evolutionarily distant protein templates, with the first identification and molecular characterization of an ALS targeting herbicide protective mutant in *Juncus effusus*. The efficacy of the novel protein-template transcript assembly methodology is demonstrated by adding gymnosperms and Polypodiopsida to the land plant backbone phylogeny. All transcript assemblies for seven species were confirmed using a novel amplicons concatenation strategy followed by Oxford Nanopore Technologies (ONT) MinION sequencing, yielding one of the most cost-effective strategies for the verification of template- assembled transcripts.

## Materials and Methods

### Plant materials

*Juncus effusus* and *Pteridium aquilinum* were sourced from a field north of Nefyn, Gwynedd, North Wales that had been drained, was topped each summer and where the rushes were spot sprayed with Imazapic and the bracken was spot sprayed with glyphosate. *Equisitum arvense* was sourced from a nearby sandy wall. *Taxus baccata* and *Pinus sylvestris* were sampled from the neighbourhood of the Boduan church, Gwynedd, North Wales. *Azolla filliculoides*, an introduced species in the UK, was collected from Nant Gwrtheyrn, Gwynedd, North Wales. Young *Ginkgo biloba* leaves were kindly donated by the Cambridge Botanical Garden, England. Where possible, plants were sampled non- destructively. For *Juncus effusus*, *Pteridium aquilinum* and *Azolla filiculoides*, to aid the discovery of herbicide resistant mutations, six individual plants were analysed (combined root, stem and leaf tissues). *Berberis thunbergii* leaves were collected from a plant in Wickham Park, Manchester, UK and *Magnolia sinostellata* leaves were taken from a shrub purchased from Thompson and Morgan, Ipswich, Suffolk, UK.

#### Choice of Genes

Initial gene lists for low copy number genes were obtained from Zeng et al. (2014). Their longest transcripts were chosen in preference and the Arabidopsis identifier for this was taken from the publication. This was used to search for a *Selaginella moellendorfii* gene orthologue using the Ensembl plants genome browser (Kersey et al. 2018). If no *Selaginella* protein could be found, the transcript was excluded from further analyses. The *Selaginella* protein was employed to find the closest orthologue in *Azolla filliculoides* using FernBase (https://www.fernbase.org/). The *Amborella trichopoda* and *Sorghum bicolor* orthologues were identified from Ensembl’s orthology (Compara) interface (Vilella et al. 2009). *Ananas comosus* and *Phoenix dactylifera* orthologues were found from NCBI’s protein blast (Johnson et al. 2008). A list of all orthologues identified from sequenced genomes is provided as Online Resource 1. For the gymnosperms (naked seed plants) few complete protein entries were found in GenBank. As a result, NCBI’s transcript sequence assembly (TSA: https://www.ncbi.nlm.nih.gov/genbank/tsa/) database was mined for full-length transcripts, using the fern and *Amborella* assemblies as reference sequences. One or two of the best-hit sequences were chosen for each of the 14 low copy number transcripts. These were translated to protein, which were added to the protein template file. A similar process was used to identify herbicide target transcripts (except that the condition that the transcript had to be present in *Selaginella* was dropped). Transcript sequences were concatenated into a single file for further use.

#### Transcript assembly and Primer Design

Transcriptomic short read datasets for species of interest were extracted from NCBI’s sequence read archive (SRA) (see Online Resource 2 for details). For the protein coding genes, herbicide targets (*ALS*, *EPSPS*, *GS*, *ACCase*, *HPPD*, *PDS* and *PPO*) as well as the low copy number genes (*SMC1*, *SMC2*, *MCM5*, *TIN1*, *NM1*, *FMNLO*, *MED17*, *EYE*, *RabGAP*, *FAAH*, *ABC1K3*, *TIF2*, *TDN* and *ALS*) the reference proteins as chosen above (see Supplementary Electronic Materials 2 for details) were used. The concatenated FASTA protein file was indexed by Diamond (Buchfink et al. 2015). Diamond was then employed to map the SRA files (converted to FASTA using a custom script) to the reference proteins. Left and right reads were mapped separately and mapped reads were concatenated into a read identifier file. The reads matching the protein of interest were extracted with Seqtk (https://github.com/lh3/seqtk) prior to assembly with SPAdes (Bankevich et al. 2012) using default parameters. If the SPAdes assembly yielded a full-length transcript, assembly was stopped at this stage. However, if only a partial transcript was used, the transcript was employed to extract more reads using mirabait (Chevreux et al. 1999) with a k-mer of 29. These reads were then assembled with SPAdes. Repeated rounds of bait and assembly were conducted until a full-length transcript was obtained.

Using complete transcript sequences, primers for ONT sequencing were designed using DegePrime (Hugerth et al. 2014) or NCBI’s primer design tool (https://www.ncbi.nlm.nih.gov/tools/primer-blast/). Primers were designed with a Tm of 60±9°C and were placed maximally at the 5’ and 3’ ends of assembled transcripts. As a result, primers were designed to amplify the maximum length of the transcript, rather than being designed to be optimal. Due to this constraint, many primers had to be designed manually. Primers sequences are provided in Supplementary Electronic Materials 3. For MinION sequencing, primers were designed with two extensions, a 5bp barcode sequence specific to the individual species+transcript amplicons and an extension including an unique rare cutter site (ie one not present in the two fragments to be ligated) (Fig. 1) so that specific amplicons could be co-ligated. As MinION is optimal (in terms of cost) for longer amplicons, sequences were ligated to give a total length of 50kbp before sequencing.

**Fig. 1.**
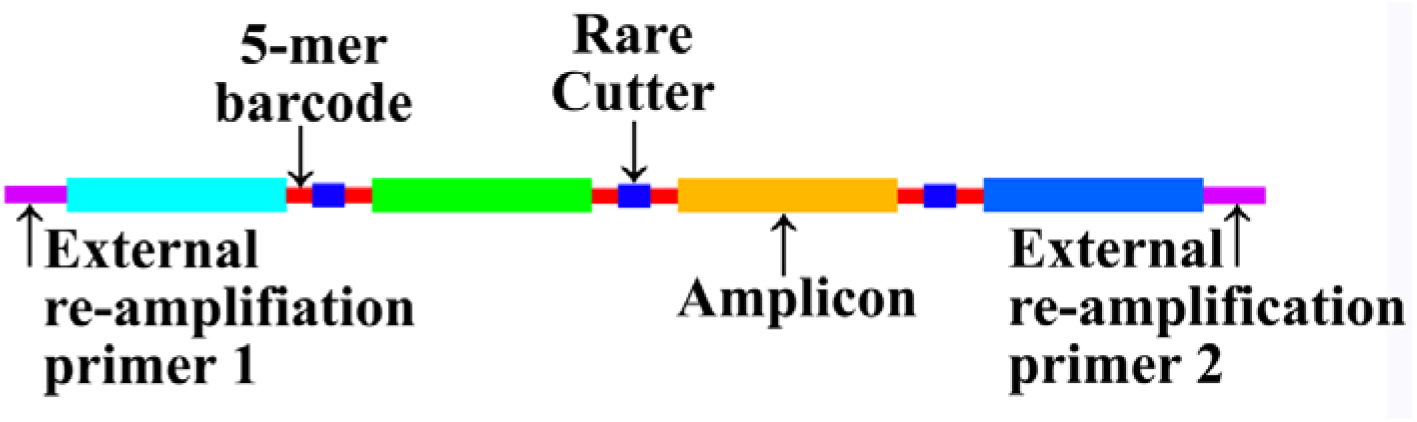
Schematic for the protein-based assembly approach Schematic representation of the Diamond (Buchfink et al. 2015) based workflow employed to assemble transcripts from raw RNA-seq reads employed in this project. Protein templates are used as the basis for transcript assembly, rather than the more usual genomic or transcript templates from closely related species. This approach effectively eliminates the evolutionary distance barrier encountered in traditional reference-based assembly techniques.

#### cDNA Library Preparation

Wherever possible young tissue was sourced and snap frozen in liquid nitrogen prior to RNA extraction. Leaf tissue (5g) was ground in a pestle and mortar under liquid nitrogen and ground tissue was immediately utilized for RNA extraction using the CTAB and activated charcoal methodology (Rajakani et al. 2013) prior to purifying mRNA using Oligo- dT cellulose (Sigma Aldrich, UK). cDNA was prepared using the ThermoFisher Scientific (UK) High-Capacity RNA-to-cDNA kit using the kit protocol.

#### MinION Sequencing and Sequence Assembly

Amplification, column purification, ligation and Oxford Nanopore Technologies sequencing were performed by CCS, Waterbeach, Cambridge, UK. For MinION sequencing, concatenated amplicons were barcoded and sample preparation was carried out using the Genomic DNA Sequencing Kit SQK-MAP-006 (Oxford Nanopore Technologies) following the manufacturer’s instructions, including the optional NEBNext FFPE DNA repair step (NEB). A 6μl aliquot of pre-sequencing mix was combined with 4 μl Fuel Mix (Oxford Nanopore), 75μl running buffer (Oxford Nanopore) and 66μl water and added to the flow cell. The 48h genomic DNA sequencing script was run in MinKNOW V0.50.2.15 using the 006 workflow. Metrichor V2.33.1 was used for base calling. Raw data from a single MinION flow cell was initially analysed using the Oxford Nanopore base calling software and binned into ‘pass’ or ‘fail’ based on a threshold set at approximately 85 % accuracy (Q9). In our hands, a 12-hour MinION run yielded 90% of the highest quality and runs were terminated after 12 hours. Reads were separated according to barcode and barcode sequences were removed. Each amplicon was assembled individually with Canu (Koren et al. 2017), using Canu’s in- built error correction. All MinION derived sequences: 94 low copy number transcripts and 32 herbicide target transcripts from this study have been deposited in ENA under the project accession PRJEB30976.

#### Sequence Alignment and Analysis

*ABC1K3, ALS, EYE, FAAH, FMNLO, MCM5, MED17, NM1, RabGAP, SMC1, SMC2, TIF2, TIN1* and *TDN* sequences were oriented in the correct strand direction using Blast (ref). Each individual gene was taken and combined with additional sequences across the main genera of vascular land plants (see the associated Supplementary Electronic Materials). Sequences were aligned with SATÉ (Liu et al. 2009) and alignments were optimized with Prank (Lo□ytynoja et al. 2012), an indel-aware alignment optimizer.

Alignments were trimmed to remove long 5’ and 3’ overhangs and missing sequence was padded with Ns. Gaps of >20aa induced in the alignment by a single species were trimmed to 20aa. Individual alignments were merged with a custom PERL/Shell script set prior to phylogenetic analyses. For mutation analyses, transcriptomic sequences for the seven herbicide target genes were converted to protein with the Expasy translate tool (https://web.expasy.org/translate/) and the protein sequences were aligned with the Stretcher Needleman-Wunsch aligner of the EMBOSS package (Rice et al. 2000).

#### Phylogenetic Analyses

Previous analysis (D. Lloyd Evans, in preparation) indicated that acetolactate synthase provided a phylogenetic signal that congruent with species evolution. As a result, ALS was added to the list of low copy number genes.

The concatenated abc1k3, als, eye, faah, fmnlo, mcm5, nm1, rabgap, smc1, smc2, tif2, tin1 and tdn assemblies were employed for phylogenetic analyses. The low copy number transcripts were divided into their five component partitions. Each partition was tested with IQ-Tree’s (Nguyen et al. 2015) model testing using the – TESTNEWONLY (including rate heterogeneity) switch and the AICc criterion to determine the best-fit evolutionary models. The optimal models were as follows: abc1k3: GTR+F+R5; als: TVMe+R5; eye: TVM+F+R5; faah: GTR+F+R4; fmnlo: GTR+F+R3; mcm5: TIM2+F+R5; med17: GTR+F+R6; nm1: GTR+F+R5; rabgap: GTR+F+R5; smc1: GTR+F+R5; smc2: GTR+F+R4; tif2: GTR+F+R4; tin1: TVM+F+R5; tdn: GTR+F+R5. The partitions determined above and their closest model equivalents were used for all subsequent analyses. Bayesian analyses were run with MrBayes (version 3.1.2) (Ronquist and Huelsenbeck 2003), Maximum Likelihood analyses and SH- aLRT single-branch analyses were run with IQ-Tree (Nguyen et al. 2015).

Bayesian Markov Chain Monte Carlo (MCMC) analyses were run on the partitioned alignment, with partitions unlinked, using MrBayes 3.1.2 (Ronquist and Huelsenbeck 2003), using four chains (3 heated and 1 cold) with default priors run for 20 000 000 generations with sampling every 100th tree. Two independent MrBayes analyses, each of two independent runs, were conducted. To avoid any potential over-partitioning of the data, the posterior distributions and associated parameter variables were monitored for each partition using Tracer v 1.6 (Rambaut et al. 2017). High variance and low effective sample sizes were used as signatures of over- sampling. Burn-in was determined by topological convergence and was judged to be sufficient when the average standard deviation of split frequencies was <0.001 along with the use of the Cumulative and Compare functions of AWTY (Nylander et al. 2008). The first 50 000 (25%) sampled generations were discarded as burn-in, and the resultant tree samples were mapped onto the reference phylogram (as determined by maximum likelihood analysis) with the SumTrees 4.0.0 script of the Dendropy 4.0.2 package (Sukumaran and Holder 2010).

To provide support for the Maximum Likelihood phylogeny, a total of 10 000 bootstrap replicates were analysed with IQ-Tree (Nguyen et al. 2015). Replicate trees were summarized internally within the IQ-Tree package. As an additional measure of branch confidence, SH- aLRT analyses were run for 10000 replicates with IQ-TREE, using the -bnni option (Nguyen et al. 2015) to reduce the risk of overestimating branch supports due to severe model violations. The final concatenated alignment (with partitioning schema) and reference phylogeny have been submitted to TreeBase under the accession TB2:S23996.

#### Structural Modelling

Modelling of *Juncus effusus* ALS protein was initially performed on the Phyre^2^ server (Kelley et al. 2015) by mapping the *Juncus effusus* ALS protein sequence to the known crystal structure of *Arabidopsis* ALS. Subsequently, both *Juncus effusus* ALS models were threaded onto our previous model of sugarcane ALS complexed with imazapyr (Lloyd Evans and Joshi 2016b; Koetle et al. 2018) using UCSC Chimera (Pettersen et al. 2004) prior to molecular dynamics simulation.

The resulting structure was prepared for molecular dynamics (MD) simulation using the Protein Preparation Wizard of the Maestro molecular modelling software (v.9.6; Schro□dinger, Inc.). The model included all hydrogen atoms from the start, but the polar interactions of the *His* residues were manually checked and the protonation states selected to optimize the hydrogen bond network.

MD simulations were performed to confirm that the 3D structure was stable without unfolding or any significant changes in secondary structure. The Gronigen Machine for Chemical Simulations (GROMACS) (Abraham et al. 2015) with the CHARMM force-field was employed for this purpose and solvated our model in a cubic box with TIP3P water. The system was charge equilibrated with 6 sodium ions before being energy minimized. After energy minimization, the systems were equilibrated by position restrained molecular dynamics at constant temperature of 300 K and a constant pressure of 1 atm for about 100 ps before running a 200 ns molecular dynamics simulation using the CHARMM force-field. The final model was compared with both the original threaded model and the *Arabidopsis* ALS crystal structure to ensure conformational stability.

#### Mutation modelling

The mutation site identified in *Juncus effusus* ALS was analyzed in the final docked structure. For site directed mutagenesis, FoldX 3.04b (Schymkowitz et al. 2005) was employed. The two mutations were engineered independently and FoldX was employed to minimize the structure and to predict the effects of the mutation on protein stability. As FoldX removes heteroatoms from its models, the cofactors and herbicide were re-docked with the molecules (as above) prior to functional analyses.

## Results

### Transcript Assembly

Based on the set of proteins described by Zhang et al. (2012) and Zeng et al. (2014) the *Arabidopsis thaliana* gene identifier was taken from the papers and input into Ensembl Plants (http://plants.ensembl.org). Where orthologues existed in *Selaginella moellendorfii* and *Physcomitrella patens* (see Supplementary Electronic Materials 1) these were taken as potential candidate genes for land plant universality.

The main target species for this study were *Juncus effusus, Equisetum arvense*, *Pteridium aquilinum and Azolla filliculoides*, which are orphan weeds in crops, particularly pastureland or are potentially invasive species. They are evolutionarily distant (at least 100 million years) from the closest assembled species. This means that the usual mapping assembly methods will not work on these species. Instead, a new methodology was developed, based on read mapping to protein references, as detailed in Fig. 2.

**Fig. 2.**
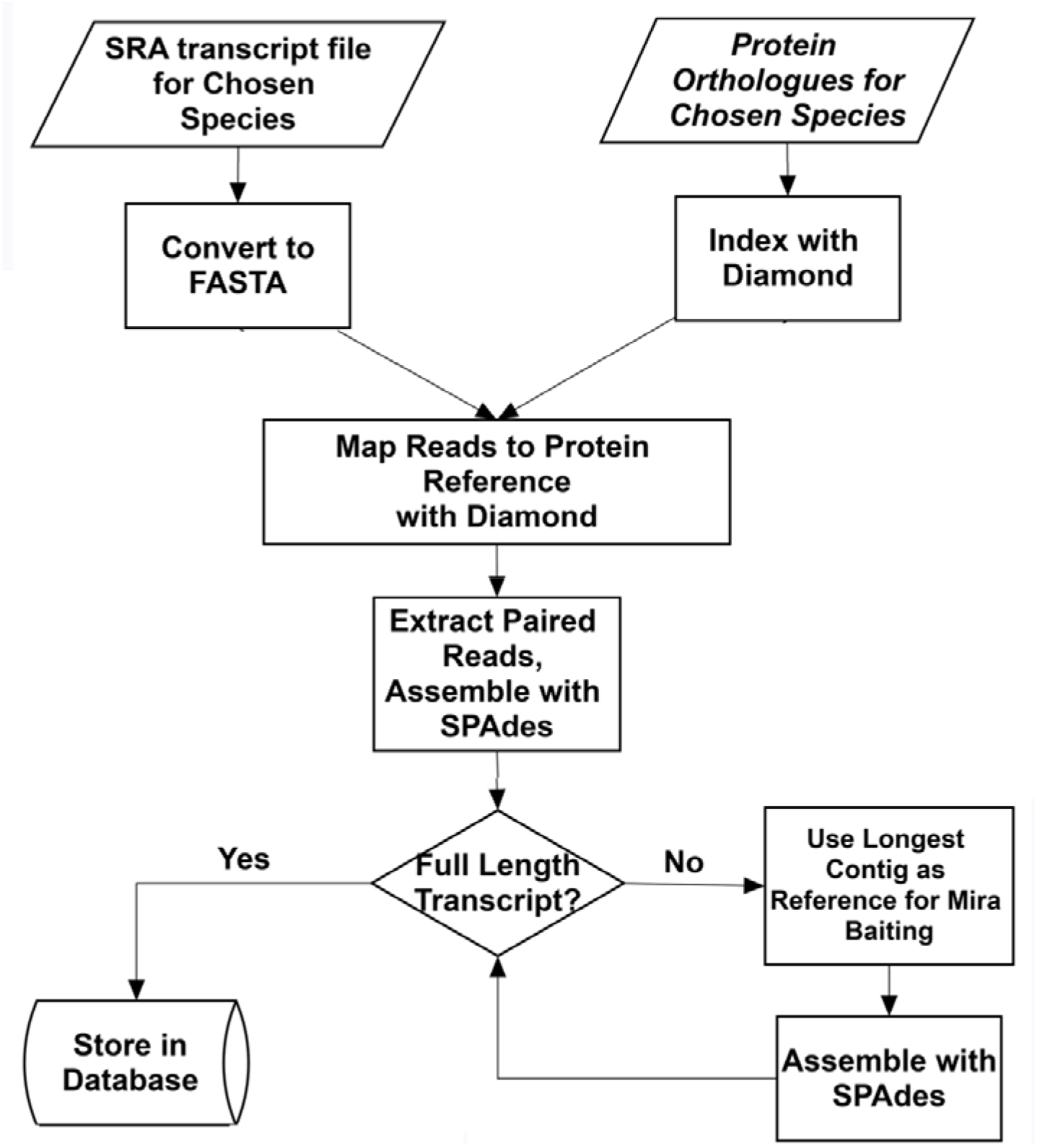
Workflow for the transcriptome assembly process employed in this study Schematic detailing the novel protein-based transcriptome methodology employed in this study. Representative proteins related to the transcript to be assembled are chosen, these are used to select a subset of related reads from the total read pool using Diamond. These reads are assembled with SPAdes. If the assembly is full length, the pipeline is terminated. If the assembly is not full length, the assembled transcript is uses as bait for mirabait to re-select reads from the read pool. These reads are then re-assembly. Assembly continues until no more reads can be extracted or a full-length coding sequence is obtained.

Previously, Zhang et al. (2012) and Zeng et al. (2014) employed a PCR-based approach using universal primers was used to assemble fragments of these proteins. In the novel methodology presented in this paper, proteins were used to bait raw reads from read pools published in SRA. The transcripts were assembled, extended if possible, and the longest assembly was used for further studies (the aim was to obtain full-length transcripts for each gene). For the low copy number genes, as well as being present in plants from bryophytes to crown monocots and dicots, the genes had to be present in species that were both antecedent and descendant of the species of interest. Table 1 lists the 14 transcripts chosen for this study, including Gene identifier, gene name, identifier for the *Arabidopsis* reference gene and the identifier of the *Sorghum bicolor* orthologue.

**Table 1.** Key to the genes/transcripts employed as low copy number references in this study The table lists the gene identifiers and gene names for the 14 genes and associated transcripts employed for the low copy number phylogeny in this paper. Also given are the reference *Arabidopsis thaliana* gene identifiers and the identifiers for the orthologues of these genes in *Sorghum bicolor* (for the full list of genes and orthologues from sequenced genomes used for phylogenetics see Online Resource 2).

| **Gene ID** | **Gene Name** | **Arabidopsis Identifier** | **Sorghum orthologue** |
| --- | --- | --- | --- |
| ABC1K3 | Protein ACTIVITY OF BC1 COMPLEX KINASE 3, chloroplastic | AT1G79600 | SORBI_3009G093100 |
| ALS | Acetolactate synthase, chloroplastic | AT3G48560 | SORBI_3004G155800 |
| EYE | Embryo yellow eye protein/Conserved oligomeric Golgi complex component-related | AT5G51430 | SORBI_3010G223400 |
| FAAH | Fatty acid amide hydrolase | AT5G64440 | SORBI_3006G002400 |
| FMNLO | tRNA-dihydrouridine(47) synthase [NAD(P)(+)]-like | AT4G38890 | SORBI_3006G010400 |
| MCM5 | DNA replication licensing factor MCM5 | AT2G07690 | SORBI_3004G331600 |
| MED17 | Mediator of RNA polymerase II transcription subunit 17 | AT5G20170 | SORBI_3003G104000 |
| NM1 | Nodal Modulator 1 | AT3G62360 | SORBI_3003G143800 |
| RabGAP | RabGAP/TBC domain-containing protein | AT5G52580 | SORBI_3004G264400 |
| SMC1 | Structural maintenance of chromosomes protein 1 | AT3G54670 | SORBI_3007G161000 |
| SMC2 | Structural maintenance of chromosomes protein 2-1 | AT5G62410 | SORBI_3003G393100 |
| TIF2 | Translation initiation factor IF-2 like protein | AT4G11160 | SORBI_3006G093300 |
| TIN1 | Tunicamycin induced protein | AT5G64510 | SORBI_3003G308400 |
| TDN | Transducin/WD40 repeat-like superfamily protein | AT2G20330 | SORBI_3001G227400 |

To fill in the land plant phylogeny, the following species were also analysed: *Lygodium japonicum*, *Ophioglossum vulgatum*, *Gnetum montanum*, *Acorus calamus*, *Magnolia sinostellata*, *Chloranthus japonicus*, *Platanus occidentalis*, *Berberis thungbergii* and *Lilium longiflorum*. In all, the following transcripts were targeted: abc1k3, als, eye, faah, fmnlo, mcm5, med17, nm1, rabgap, smc1, smc2, tif2, tin1, tdn. Online Resource 3 gives the genes assembled, maximum transcript length, length of CDS (if fully assembled) and rounds of assembly performed for each species analysed. As far as possible data were extracted from whole genome assemblies (either from Ensembl (Kersey et al. 2018) or Phytozome (Goodstein et al. 2011) or the data of Zeng et al. (2014). However if the assemblies/sequenced reads from Zeng et al. were severely truncated and could not readily be re-assembled an alternate species from the same genus/family was used. If genes for a given species from the genome repositories were severely truncated or obviously misassembled, an attempt was made to locate a full-length transcript in GenBank. If no such transcript was available the transcript was assembled using the pipeline presented in Fig. 2 from RNA-seq data. Species and SRA transcriptomic files used to assemble the transcripts are detailed in Online Resource 2.

Where cognate datasets were available, transcripts built using protein templates were longer than those of Zeng et al. (2014). Where transcript shotgun assemblies (TSA: https://www.ncbi.nlm.nih.gov/genbank/tsa/) were available for comparisons (as for *Juncus effusus* PRJNA345287 and *Pteridium aquilinum* PRJNA48473) the protein-baited assemblies from this study were at least as good as TSA entries and often surpassed TSA assemblies in terms of quality and transcript length — typically joining TSA fragments into a contiguous transcript and/or extending partial genes. In almost all cases, protein-guided assemblies resulted in full-length transcripts; the exceptions being when there was insufficient transcriptomic coverage (the main example being: *Chloranthus japonicus*). In no case was it impossible to assemble a transcript (or significant portion thereof) using a protein as bait for RNA-seq reads despite the considerable evolutionary distances involved between target and template.

Overall, protein-guided transcriptomic assemblies improved significantly upon *ab initio* assemblies and also yielded longer sequences as compared with universal primer approaches. This was even the case for *Azolla filiculoides*, where a genome reference existed.

An identical methodology was employed to assemble the six main herbicide-targeting gene transcripts and two of their paralogues (eight transcripts in all).

Of the total 263 transcripts targeted for assembly in this project, only 6 (2.3%) could not be assembled in their entirety (typically missing N-terminal ends of the proteins) and in every case this was due to insufficient reads in the SRA files used for assembly. Moreover, in cases where genome assembly files for the transcript assembly were missing or wrongly assembled the protein-baiting methodology detailed herein yielded full-length transcripts (11 cases; Online Resource 2).

### ONT MinION Sequencing of Transcript Assemblies

For the *Equisetum arvense*, *Pteridium aquilinum*, *Juncus effusus*, *Ginkgo biloba*, *Taxus baccata* and *Pinus sylvestris* transcript assemblies, veracity of the assembled sequences was confirmed by ONT MinION sequencing. As MinION is more cost effective for sequencing longer sequences, primers designed for transcript amplification had 5-mer barcodes appended along with rare cutter DNAse sites that allowed for specific ligation of one amplicon to the next (Fig. 1). To ameliorate loss of amplicons product in each ligation stage complete concatenated products (~50kb) had specific primer sites at the start and end so that, if necessary, long-read high-fidelity polymerases could be used to amplify the entire construct. The entire constructs were then barcoded with MinION specific barcodes prior to sequencing. Typically, up to 70 constructs of ~50kbp could be sequenced in a single MinION run (3.5Mb DNA total). The approach is shown schematically in Raw data from the concatenated reads from a single MinION flow cell was initially analysed using the Oxford Nanopore base calling software and binned into ‘pass’ or ‘fail’ based on a threshold set at approximately 85 % accuracy (Q9) and including only 2D reads (where data are generated from both the forward and reverse DNA strands as they pass through the nanopore).

Failed reads were removed and only pass reads were taken forwards to assembly. Based on Oxford Nanopore Technologies barcodes, pass reads were binned into individual components. These were assembled and error corrected with Canu (Koren et al. 2017) using the default parameters. The Canu assembly step took less than 2 hours, but the most significant delay with MinIon is the initial base-calling step. For a 48-hour base calling run, we noted that almost 50 % of pass reads were generated in the first 6 h, almost 80 % within 9 h and 90 % within 12 h. This gave a theoretical coverage of 20×, 32× and 37×, respectively. Only 31 pass (<0.1%) reads were generated in the final 12 h of the 48 h run.

Assembling only using pass reads from the first 6 h alone led to a less accurate, fragmented assembly, but subsets of pass reads taken from the first 9 or 12 h of the run generated similar assemblies to pass data from the full 48 h run. Thus base calling for 50kbp length reads needs only 14–16 hours of base calling, making end to end assembly from amplicon to assembled reads within a 24-hour timescale feasible.

Using the internal barcodes, individual amplicons were excised from the minion assembles. These sequences are available from the ENA under project identifier: PRJEB30976. In all, 130 transcripts were assembled and sequenced for this project. A further 137 transcripts were assembled from third party data and are available as a tarball in Online Resource 4.

### Phylogenetic Analysis

The phylogenetic analysis presented in Fig. 3 is largely congruent with previous studies for seed plants, Zeng et al. (2014), ferns (Shen et al. 2017) and gymnosperms (Lu et al. 2014). However, this is the first study to use low copy number nuclear genes to create a phylogeny of all major vascular plant lineages. Not all transcripts could be assembled in their entirety, thus there were gaps (padded by Ns) in the alignment. The sister relationship of the Gnetophytes with true seed plants and of the Pteridophytes with Gnetophytes + Gymnosperms is demonstrated. The phylogeny of the gymnosperms (true seed plants) is consistent with previous analyses (Zhang et al. 2012).

**Fig. 3.**
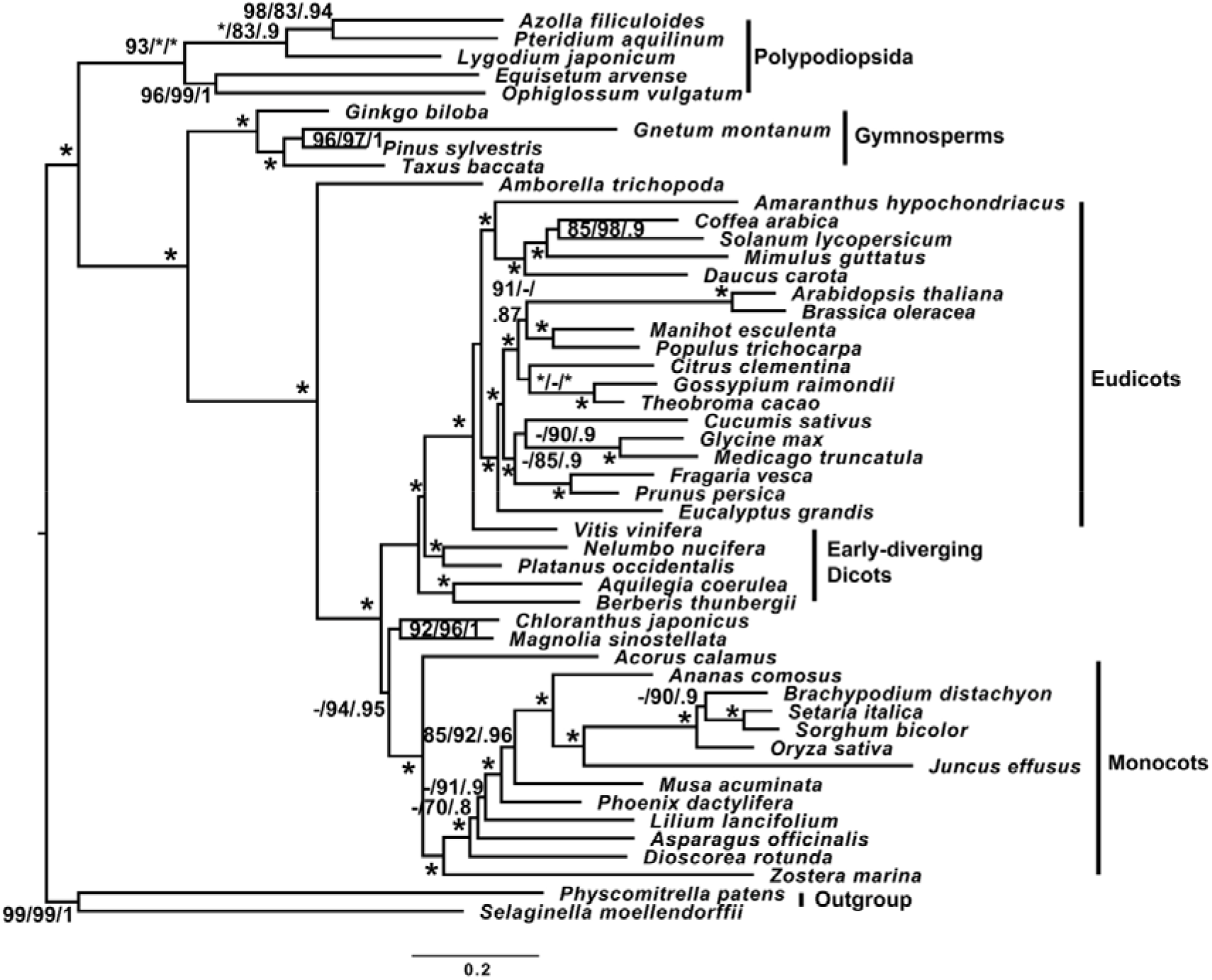
Phylogeny of vascular plants derived from 14 low copy number transcripts Phylogeny of vascular land plants derived from DNA alignments of 14 low copy number genes using *Selaginella moellendorfii* and *Physcomitrella* patens as an outgroup. Numbers next to branches represent SH-aLRT single branch test/non-parametric bootstrap and Bayesian inference. The asterisk (*) symbol represents complete support for a given branch. Only SH-aLRT values above 85%, bootstrap support above 75% and BI values above 0.7 are shown. The scale bar at the bottom represents expected numbers of substitutions per site.

### Herbicide Target Genes

All herbicide target genes for all four species of interest (*Equisetum arvense*, *Pteridium aquilinum*, *Azolla filiculoides* and *Juncus effusus*) could be assembled in their entirety using protein baiting and primers designed on the assembled sequences gave full- length transcripts. Despite sampling six plants, no variants were found in Equisetum arvense, which is not unexpected as these plants had not been subject to herbicide treatment. The same was true for *Azolla filiculoides*.

For *Pteridium aquilinum*, though treated with glyphosate no mutations were identified, however as this study only analysed RNA-datasets by PCR copy number or expression variants could not be excluded. However, in *Juncus effusus*, treated with the ALS inhibiting herbicide Imazamox, an Imidazolinone class herbicide, a single mutation was identified in one of the six plants sampled. This is an active site mutation, W574R (*Arabidopsis* numbering), W548R (*Juncus* numbering) previously described in the weed *Digitaria sanguinalis* (Li et al. 2017).

Using a previous model of sugarcane ALS complexed with Imazapyr as a template, the *Juncus effusus* ALS was modelled. The effect of the W574R mutant is shown in Fig. 4. The main effect is on Gln573 (Gln547 in *Juncus* numbering), which forms a stabilizing hydrogen bond between ALS and the herbicide. Substituting the hydrophobic side chain of Tryptophan with the positively charged Arginine induces arginine to be solvent accessible. This pulls Gln573 with it, displacing the amino acid and twisting its side chain by 6.4Å so that it can no longer physically form a stabilizing hydrogen bond with the herbicide.

**Fig. 4.**
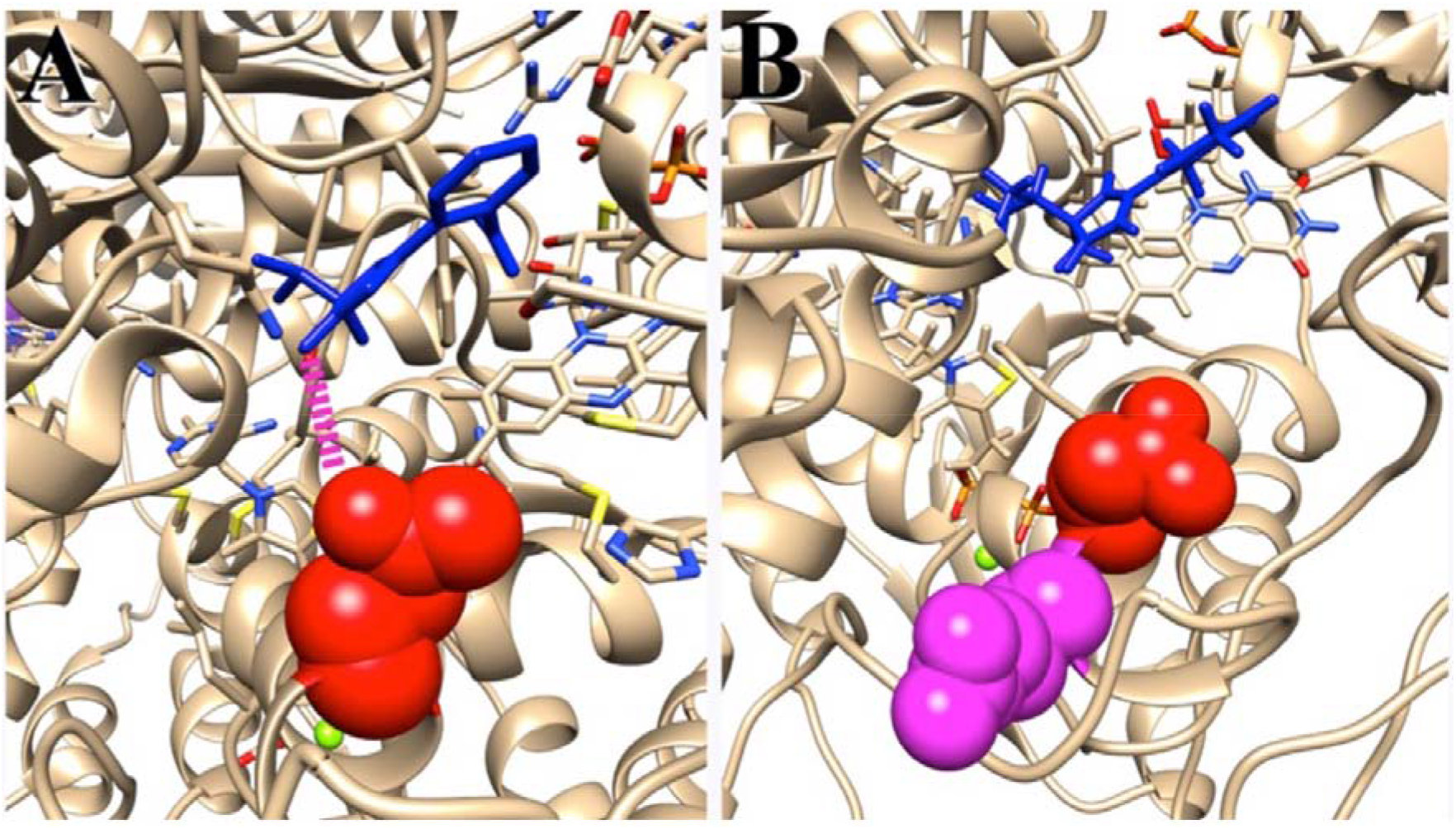
Molecular models for the active sites of wild type and mutant *Juncus effusus* acetolactate synthase (ALS) Molecular models for the active sites of acetolactate synthase in Juncus effusus showing the wild type active site (A) and the active site of the mutant form W548R (B). Both molecules have imazapyr docked in the active site (blue stick representation molecule). In part (A) Gln5437 (red space-filling representation) forms a strong stabilizing hydrogen bond (shown in pink) with the =o of the imidazoline ring of imazapyr. Mutating W548 to Arginine introduces a polar amino acid (magenta in panel (B)), which alters the conformation of Gln5437 displacing and rotating the side chain so that it can no longer form a stabilizing hydrogen bond with the herbicide. This reduces the binding affinity of the herbicide so that it can be displaced by the enzyme’s natural substrate.

## Discussion

*Juncus effusus* is a member of the Juncaceae, within the Cyperid clade of the Poales. They are sister to the Xyrids, Restids and Graminids, this latter clade containing the Poaceae or true grasses. Cyperids are over 110 million years divergent from the Poaceae and 120 million years divergent from the Bromeliaceae, which contains pineapple (*Ananas comosus*) the closest relative with an assembled genome (Bouchenak-Khelladi et al. 2014; Ming et al. 2015).

*Equisetum arvense* is a member of the Equisetaceae, the earliest diverging of the Pteridophytae. *Pteridium aquilinum* is a member of the Pteridineae and lies close to the crown ferns. Only two complete fern genomes are available, those of *Azolla filiculoides* and Salvinia cucullata, both members of the Salvinales. *Pteridium* is just over 150 million years divergent from the Salvinales and *Equisitum* is over 320 million years divergent. *Equisetum* is over 410 million years divergent from *Selaginella moellendorfii* (Shen et al. 2017).

Without a reference, transcript assembly can be highly problematic. Even with a reference as a guide, transcript assembly from short reads can suffer from issues such as read coverage, GC bias and reads shared with other transcripts. This can and will result in truncated transcripts and chimaeric transcripts. Mapping to a reference can help generate improved transcripts and will remove contamination due to bacterial, fungal and viral contaminants. Typically, genomes or reference transcriptomes are used as templates. Here, a novel methodology using protein sequences as templates is employed. For the four main species in this study, 22 transcripts each were assembled, including low copy number genes and herbicide targets. Each of the species can be considered an orphan, in a genome context, in that it has no genome assembly and is evolutionarily distant from the closest reference, meaning that DNA or RNA templates cannot be employed. Though two fern genomes, those of *Azolla filliculoides* and *Salvinia cucullata* are available, these are 335 million years divergent from Equisetum. Moreover, of the 14 low copy number genes and 8 herbicide target genes chosen for this study not all have complete CDSes in the *Azolla* and *Salvinia* genomes (http://www.fernbase.org). But, using additional transcriptomic datasets it was possible to assemble complete CDSes for each representative sequence in Azolla. Equisetum arvense is so distant from its nearest sequenced and assembled relative that standard template-based methods (mapping to a reference, bait and assemble) failed to identify sufficient reads for any of the genes for a transcript to be assembled (typically no matching reads were recovered). However, using the new protein-based read mapping methodology described in this paper, all CDSes apart from one, that of tdn (transducing) could be assembled in their entirety and this assembly failure was not a result of the methodology employed, rather there were insufficient reads to fully assemble the transcripts. The Equisetum arvense transcripts were translated and added to the protein file. The extended file was used as a baitset and employed to assemble the transcripts from the next fern species, Pteridium aquilinum. In this case all transcripts except for med17 assembled in their entirety (med17 failed for the same reason as above). These transcripts were translated and the proteins were added to the bait file. The new protein set was used to assemble the low copy number proteins from *Ophiglossum vulgatum*, were all transcripts assembled completely. In each case the extended protein set was used to bait reads from the next species. When all transcripts from all species had been assembled, the complete protein set was employed to see if the partially assembled transcripts could be finished in their entirety.

For the gymnosperm species analysed in this study, a similar process was employed. Proteins from *Amborella trichopoda*, *Selaginella moellendorfii* and *Ophioglossum vulgatum* were used as a baitset. Using these proteins, the complete low copy number transcript set from *Pinus sylvestris* was assembled in its entirety. This protein set was added to the total protein set and the extended set was used to bait reads from *Taxus baccata*. The assembled transcripts were translated and the proteins were added to the protein set then the next most evolutionarily distant species had its transcript set assembled. This was continued until all the target species from the gymnosperms had their low copy number transcripts assembled. For *Juncus effusus*, proteins from *Setaria italica*, *Brachypodium distachyon*, *Ananas comosus*, *Phoenix dactylifera* and *Musa acuminata* were employed as the bait set. For all these species, primers spanning the CDSes were designed and the full length CDS was sequenced with ONT MinION. For the other species in the phylogeny, where no full length CDS was present in GenBank and no genome reference existed the absent or truncated transcript was assembled from RNA-seq datasets using a combination of standard bait and assembly or protein-directed bait an assembly. In all, 256 transcripts were assembled for this study, of which 250 assembled in their entirety and 130 of these were sequenced to confirm the assembly process. Most assembly problems were encountered with *Chloranthus japonicus*, simply because the RNA-seq data available were of insufficient depth to yield full-length transcripts for four genes. However, in every case it was possible to extend the transcripts over and above the length of those currently deposited in GenBank. For sequenced genomes, if there were problems with the transcripts these were re-assembled, yielding 11 additional transcripts. All these are detailed in Online Resource 2.

### Herbicide target genes

For the herbicide target genes analysed in this study, from three fern species and one member of the Juncaceae this is the first time that a directed assembly and sequencing analysis has been performed for any of these species or groups. The species were chosen due to their evolutionary divergence from know herbicide targets and because all the species are either weeds or are invasives in the UK. Of the eight genes chosen all could be assembled in their entirety using protein-baiting of RNA-seqs and all sequences were confirmed by ONT MinION sequencing. In each case six distinct individuals were sequenced separately. Though sequences were obtained for each isolate, a mutation was only discovered in *Juncus effucus* acetolactate synthase. This variant is cognate to the W574R (Arabidopsis numbering) mutation previously described in *Digitaria sanguinalis* (Li et al. 2017). However, this is the first report of the sequence of this transcript and the first discovery of a herbicide inhibiting mutant in any Poales species. This is also the first report of a herbicide-associated mutation in response to the herbicide Imazamox. Though the mutation is previously known in other plant species, its precise mechanism of action had not been determined. As a result, based on a previously generated model of the active sugarcane ALS dimer complexed with imazapyr, a model of the functional dimer of *Juncus effusus* was generated by threading. In this model (Fig. 4) Gln573 forms a strong hydrogen bond with the =O side chain of the imidazole ring in imazapyr (and imazamox in which the imidazole ring is identical) stabilizing the herbicide in the active site. Mutating Tryptophan 574 to an Arginine transforms an amino acid with a hydrophobic side chain into one with a positively charged side chain. Arginine prefers to be solvent exposed, whilst Tryptophan prefers to be buried. As a result of the mutation, the Arginine side-change emerges from the molecule to be solvent exposed. This pulls the Glutamine residue along with it so that Gln573 can no longer form a stabilizing hydrophobic bond with the herbicide. This destabilizes the herbicide interaction in the active site and allows the native substrate of ALS to displace the herbicide. Thus, the functional effect of the W574R herbicide-binding mutant is explained. For the other herbicide targets assembled and sequenced these are the first reports of these transcripts in any of the four species analysed. *Equisetum arvense* is particularly interesting as it is resistant to many of the commonly applied herbicides. Though no obvious protective mutations could be detected in the sequenced genes, this does not preclude the effects of gene copy numbers and up-regulation of expression. The sequences derived for this paper will allow for future studies on the mechanisms of herbicide tolerance/resistance in *Equisetum arvense*.

For bracken, *Pteridium aquilinum*, which is potentially deleterious to human health, the herbicide target transcripts obtained in this study can be employed as references for herbicide development and to test for potential copy number variants and novel herbicide binding mutations.

### Plant Phylogenetics

Previous large-scale analyses employing low copy number genes to analyze angiosperms as a whole (Zhang et al. 2012; Zeng et al. 2014) have used five or 59 low copy number genes, respectively. In this study, 14 low copy number genes were employed, including 13 transcripts derived from the work of Zeng et al. (2014), and acetolactate synthase (ALS). Unlike the work of Zeng et al. (2014), however, full-length transcripts were used in this study. The 14 transcripts were concatenated, but were partitioned and analysed individually. The phylogram generated from the amino acid matrix is shown in Fig. 3. Overall, branch supports are good (with a few exceptions) and the tree is largely congruent with previous studies. The phylogeny is entirely consistent with that presented by Zhang et al. (2012), but with the inclusion of polypodiopsida and gymnosperns. As in the study of Zhang et al. (2012), *Magnolia* is sister to *Chloranthus* and both are sister to the monocots. Whilst the sister relationship between Magnolia and Chloranthus has good support, the relationship between these two genera and the Monocots is not as well supported (Fig. 3). Indeed, studies with larger numbers (59) of transcripts that include introns (Zeng et al. 2014) or with very large numbers of genes/transcripts (Morris et al. 2018) place Chloroanthus as sister to the Eudicots whilst Magnolia is sister to the Eudicots + Chloranthus.

It should be noted, however, the supplementary data presented by Zeng et al. (2014) based on 83 plastid genes place Chloranthus as sister to the Magnoliids with this group being sister to the Eudicots + Monocots. Such divergence in phylogenetic signal between different datasets is consistent with network (reticulate) evolution (Linder and Rieseberg 2004) between the magnoliids, chloranthaceae, eudicots and monocots and resolution of the evolutionary history between these groups of true seed plants may require considerably more work.

However, the phylogeny described in this paper places *Amborella* as sister to the Eudicots and Monocots (supporting previous work) with the Gymnosperms as sister to (Amborella + Eudicots + Monocots) and polypodiopsida as sister to Gymnosperms + (Amborella + Eudicots + Monocots). This means that, broadly, the fourteen transcripts chosen herein can be employed to determine the phylogenetics of all vascular land plants and the methodology developed places the sequencing of select genes from orphan plants within the grasp of most groups (sequencing costs being in the order of < $1000).

## Conclusion

A novel method for assembling transcripts from RNA-seq data using proteins as a baiting step is introduced. This overcomes the ~20 million-year (30% sequence divergence) barrier at which template-based assembly is currently impossible. This is allied with a new methodology for rapidly, efficiently and cost-effectively confirming these assemblies by direct sequencing with ONT MinION technology. The methodology is confirmed by assembling and sequencing eight herbicide target genes from ferns and *Juncus effusus* as well as assembling 261 low copy number transcripts (14 for each species) and sequencing 150 of these transcripts. The utility of these transcripts for phylogenetic studies across all vascular land plant was examined. Phylogenetic studies demonstrated that the transcripts chosen were effective, yielding robust phylogenies that included Gymnosperms and Polypodiopsida for the first time. This means that a cheap and effective technique now exists for analysing the phylogenetics of land plants with a consistent set of transcripts. This allows existing RNA-seq datasets to be mined for phylogenetically informative data.

For the first time, the six main known herbicide targets from three fern and one Poaceae species are revealed — all of these species being weeds or invasives. This provides a strong basis for future work on these species. Moreover, for the first time, a herbicide mutant is described for a member of the Juncaceae (the first in non-grass Poaceae) and the molecular mode of action of this mutation is elucidated and described.

## Compliance with Ethical Standards Funding

This project received no formal funding; however we are grateful to Oxford Nanopore Technologies for their support through their community access programme.

## Data Availability

All transcripts assembled in this study have been submitted to the ENA under the project identifier: PRJEB30976. Final alignments and phylogenetic trees were submitted to TreeBase: TB2:S23996. All assemblies of third-party data are available as Electronic Supplementary Material 4. Computer code developed for this project is available from GitHub: https://github.com/gwydion1/bifo-scripts.git.

## Supplementary Files

### Online Resource 1

#### ESM_1.xlsx

List of reference genes in Arabidopsis thaliana for the 14 low copy number transcripts analysed in the main manuscript. Orthologues from ENSEMBL Plants or Phytozome are given for 33 species whose complete genomes have been sequenced. Blue cells represent genes that were absent or misassembled in the genomes.

### Online Resource 2

#### ESM_2.xlsx

List of species for which the 14 low copy number genes and 7 herbicide target genes were assembled including a list of all SRA files used in assemblies. Yellow cells represent transcripts that could only be assembled in part.

### Online Resource 3

#### ESM_3.xlsx

List of all species for which transcripts were assembled in this study. Also given are the length of the transcripts and length of coding regions within the transcripts. For those transcripts that were sequenced the amplification primers employed are also provided.

### Online Resource 4

#### ESM_4.tar.gz

Gzipped tar file containing all the transcript assemblies constructed from third party data that were not sequenced. The file is a concatenated set of folders for each species whose transcripts were assembled.

## References

Abraham MJ, Murtola T, Schulz R, Páll S, Smith JC, Hess B, Lindahl E (2015) CROMACS: High performance molecular simulations though multi-level parallelism from laptop to super computers. Software X1:19–25.

Bankevich A, Nurk S, Antipov D, Gurevich AA, Dvorkin M, Kulikov AS, Lesin VM, Nikolenko SI, Pham S, Prjibelski AD, Pyshkin AV (2012) SPAdes: a new genome assembly algorithm and its applications to single-cell sequencing. J Comput Biol 19:455–477.

Bens, M., Sahm, A., Groth, M., Jahn, N., Morhart, M., Holtze, S., Hildebrandt, T.B., Platzer, M. and Szafranski, K., 2016. FRAMA: from RNA-seq data to annotated mRNA assemblies. BMC Genomics 17:54.

Berlicki L (2008) Inhibitors of glutamine synthetase and their potential application in medicine. Mini reviews in medicinal chemistry 8:869–878.

Bouchenak-Khelladi Y, Muasya AM, Linder HP (2014) A revised evolutionary history of Poales: origins and diversification. Botanical Journal of the Linnean Society 175:4–16.

Buchfink B, Xie C, Huson DH (2015). Fast and sensitive protein alignment using DIAMOND. Nature methods 12:59.

Chevreux C, Wetter T, Suhai S (1999) Genome sequence assembly using trace signals and additional sequence information. In: German Conference on Bioinformatics. 99:45–56.

Clauson□Kaas F, Jensen PH, Jacobsen OS, Juhler RK, Hansen HCB (2014) The naturally occurring carcinogen ptaquiloside is present in groundwater below bracken vegetation. Environmental toxicology and chemistry 33:1030–1034.

Curran W et al. Eds (2000) Weed Control Manual 2000. Meister Pub. Co. Willoughby, OH. Galpin OP, Whitaker CJ, Whitaker RH, Kassab JY (1990). Gastric cancer in Gwynedd. Possible links with bracken. British Journal of Cancer 61:737.

Garcia D, Ramos AJ, Sanchis V, Marín S (2012) Effect of *Equisetum arvense* and *Stevia rebaudiana* extracts on growth and mycotoxin production by *Aspergillus flavus* and *Fusarium verticillioides* in maize seeds as affected by water activity. International Journal of Food Microbiology 153:21–27.

Góngora-Castillo E, Buell CR (2013) Bioinformatics challenges in de novo transcriptome assembly using short read sequences in the absence of a reference genome sequence. Natural product reports 30:490–500.

Goodstein DM, Shu S, Howson R, Neupane R, Hayes RD, Fazo J, Mitros T, Dirks W, Hellsten U, Putnam N, Rokhsar DS (2011) Phytozome: a comparative platform for green plant genomics. Nucleic Acids Res 40:D1178–D1186.

Hao GF, Zuo Y, Yang SG, Yang GF (2011) Protoporphyrinogen oxidase inhibitor: an ideal target for herbicide discovery. CHIMIA International Journal for Chemistry 65:961–969.

Hasebe M (1999) Evolution of reproductive organs in land plants. Journal of Plant Research 112:463–474.

Hill RJ, Foland D (1986) Equisetum. In: Poisonous Plants of Pennsylvania. Dept. of Agric., Bureau of Plant Industry. Harrisburg, PA. pp 67–68.

Hirono I, Ito M, Yagyu S, Haga M, Wakamatsu K, Kishikawa T, Nishikawa O, Yamada K, Ojika M, Kigoshi H (1993). Reproduction of progressive retinal degeneration (bright blindness) in sheep by administration of ptaquiloside contained in bracken. Journal of Veterinary Medical Science, 55:979–983.

Huerta-Cepas J, Szklarczyk D, Forslund K, Cook H, Heller D, Walter MC, Rattei T, Mende DR, Sunagawa S, Kuhn M, Jensen LJ (2015) eggNOG 4.5: a hierarchical orthology framework with improved functional annotations for eukaryotic, prokaryotic and viral sequences. Nucleic Acids Research 44:D286–D293.

Hugerth LW, Wefer HA, Lundin S, Jacobsson HE, Lindberg M, Rodin S, Engstrand L, Andersson AF (2014) DegePrime: a program for degenerate primer design for broad taxonomic-range PCR for microbial ecology studies. Appl. Environ. Microbiol. 80:5116–5123.

Johnson M, Zaretskaya I, Raytselis Y, Merezhuk Y, McGinnis S, Madden TL (2008) NCBI BLAST: a better web interface. Nucleic Acids Research 36:W5–W9.

Jones E, Simpson DA, Hodkinson TR, Chase MW, Parnell JA (2007) The Juncaceae- Cyperaceae interface: a combined plastid sequence analysis. Aliso: A Journal of Systematic and Evolutionary Botany 23:55–61.

Kelley LA, Mezulis S, Yates CM, Wass MN, Sternberg MJ (2015) The Phyre2 web portal for protein modeling, prediction and analysis. Nature Protocols, 10:845.

Kersey PJ, Allen JE, Allot A, Barba M, Boddu S, Bolt BJ, Carvalho-Silva D, Christensen M, Davis P, Grabmueller C Kumar N (2017). Ensembl Genomes 2018: an integrated omics infrastructure for non-vertebrate species. Nucleic Acids Res 46:D802–D808.

Koetle MJ, Lloyd Evans D, Singh V, Snyman SJ, Rutherford RS, Watt MP (2018) Agronomic evaluation and molecular characterisation of the acetolactate synthase gene in imazapyr tolerant sugarcane (*Saccharum* hybrid) genotypes. Plant cell reports 37:1201–1213.

Kukorelli G, Reisinger P, Pinke G (2013) ACCase inhibitor herbicides–selectivity, weed resistance and fitness cost: a review. Int. J. Pest Management 59:165–173.

Kraehmer H, van Almsick A, Beffa R, Dietrich H, Eckes P, Hacker E, Hain R, Strek HJ, Stuebler H, Willms L (2014) Herbicides as weed control agents–state of the art. II. Recent achievements. Plant Physiology 166:1132–1148.

Li FW, Brouwer P, Carretero-Paulet L, Cheng S, De Vries J, Delaux PM, Eily A, Koppers N, Kuo LY, Li Z, Simenc M (2018) Fern genomes elucidate land plant evolution and cyanobacterial symbioses. Nature plants 4:460.

Li J, Li M, Gao X, Fang F (2017) A novel amino acid substitution Trp574Arg in acetolactate synthase (ALS) confers broad resistance to ALS□inhibiting herbicides in crabgrass (*Digitaria sanguinalis*). Pest management science 73:2538–2543.

Liu K, Raghavan S, Nelesen S, Linder CR, Warnow T (2009) Rapid and accurate large-scale coestimation of sequence alignments and phylogenetic trees. Science 324:1561–4.

Linder CR, Rieseberg LH (2004) Reconstructing patterns of reticulate evolution in plants. Am J Bot 91:1700–1708.

Lipecki J (2006) Weeds in orchards — pros and contras. Journal of fruit and ornamental plant research 14:13.

Lloyd Evans D, Joshi SV (2016a) Complete chloroplast genomes of *Saccharum spontaneum*, *Saccharum officinarum* and *Miscanthus floridulus* (Panicoideae: Andropogoneae) reveal the plastid view on sugarcane origins. Systematics and biodiversity 14:548–571.

Lloyd Evans D, Joshi SV (2016b) Elucidating modes of activation and herbicide resistance by sequence assembly and molecular modelling of the Acetolactate synthase complex in sugarcane. Journal of theoretical biology 407:184–197.

Lloyd Evans, D. and Joshi, S.V., 2017. Herbicide targets and detoxification proteins in sugarcane: from gene assembly to structure modelling. Genome 60:601–617.

Lloyd Evans D, Joshi SV, Wang J (2019) Whole chloroplast genome and gene locus phylogenies reveal the taxonomic placement and relationship of *Tripidium* (Panicoideae: Andropogoneae) to sugarcane. BMC Evolutionary Biology 19:33.

Lo□ytynoja A, Vilella AJ, Goldman N (2012) Accurate extension of multiple sequence alignments using a phylogeny-aware graph algorithm. Bioinformatics 28:1684–91.

Lu Y, Ran JH, Guo DM, Yang ZY, Wang XQ (2014). Phylogeny and divergence times of gymnosperms inferred from single-copy nuclear genes. PloS One, 9:p.e107679.

Marshall G (1986) Growth and development of field horsetail (Equisetum arvense L.). Weed Science 34:271–275.

McCorry MJ, Renou F (2003) Ecology and management of Juncus effusus (soft rush) on cutaway peatlands. BOGFOR Research Programme. https://www.researchgate.net/publication/253779950_Ecology_and_management_of_Juncus_effusus_soft_rush_on_cutaway_peatlands. Accessed 20 Dec 2018.

Merchant S, Wood DE, Salzberg SL (2014) Unexpected cross-species contamination in genome sequencing projects. PeerJ 2:p.e675.

Ming R, VanBuren R, Wai CM, Tang H, Schatz MC, Bowers JE, Lyons E, Wang ML, Chen J, Biggers E, Zhang J (2015) The pineapple genome and the evolution of CAM photosynthesis. Nature Genetics 47:1435.

Morris JL, Puttick MN, Clark JW, Edwards D, Kenrick P, Pressel S, Wellman CH, Yang Z, Schneider H, Donoghue PC (2018). The timescale of early land plant evolution. Proc Natl Acad Sci USA 115:E2274–E2283.

Nguyen L-T, Schmidt HA, von Haeseler A, Minh QA (2015) IQ-TREE: A Fast and Effective Stochastic Algorithm for Estimating Maximum-Likelihood Phylogenies, Mol Biol and Evol 32:268–274. doi:https://doi.org/10.1093/molbev/msu300.

NNSS UK (2015) Non Native Species Secretariat UK page for *Azolla filliculoides*. http://www.nonnativespecies.org/factsheet/factsheet.cfm?speciesId=451. Accessed February 9 2019.

Nylander JA, Wilgenbusch JC, Warren DL. Swofford DL (2008) AWTY (are we there yet?): a system for graphical exploration of MCMC convergence in Bayesian phylogenetics. Bioinformatics 24:581–83.

Pakeman RJ, Le Duc MG, Marrs RH (1997) Moorland vegetation succession after the control of bracken with asulam. Agriculture, ecosystems & environment 62:41–52.

Pamukcu AM, Ertu□rk E, Yalc□iner S, Milli U, Bryan GT (1978) Carcinogenic and mutagenic activities of milk from cows fed bracken fern (*Pteridium aquilinum*). Cancer Res 38:1556–1560.

Parchman TL, Geist KS, Grahnen JA, Benkman CW, Buerkle CA (2010) Transcriptome sequencing in an ecologically important tree species: assembly, annotation, and marker discovery. BMC Genomics 11:180.

Pettersen EF, Goddard TD, Huang CC, Couch GS, Greenblatt DM, Meng EC, Ferrin TE. (2004) UCSF Chimera — a visualization system for exploratory research and analysis. J Compt Chem 25:1605–1612

Pollard MO, Gurdasani D, Mentzer AJ, Porter T, Sandhu MS (2018) Long reads: their purpose and place. Human Molecular Genetics. 27:R234–R241. https://doi.org/10.1093/hmg/ddy177

Rajakani R, Narnoliya L, Sangwan NS, Sangwan RS, Gupta V (2013) Activated charcoal-mediated RNA extraction method for *Azadirachta indica* and plants highly rich in polyphenolics, polysaccharides and other complex secondary compounds. BMC research notes 6:125.

Rambaut A, Suchard MA, Xie D, Drummond AJ. Tracer, Version 1.5. http://tree.bio.ed.ac.uk/software/tracer/. Accessed 07 January 2019.

Rasmussen LH, Hansen HCB, Lauren D (2005). Sorption, degradation and mobility of ptaquiloside, a carcinogenic Bracken (*Pteridium* sp.) constituent, in the soil environment. Chemosphere 58:823–835.

Rice P, Longden I, Bleasby A (2000) EMBOSS: the European molecular biology open software suite. Trends in genetics 16:276–277.

Rodwell JS (1991) British plant communities (Vol. 2). Cambridge University Press, Cambridge, UK

Ronquist F, Huelsenbeck JP. MrBayes 3 (2003) Bayesian phylogenetic inference under mixed models. Bioinformatics 19:1572–4

Sandmann G, Schmidt A, Linden H, Bo□ger P (1991) Phytoene desaturase, the essential target for bleaching herbicides. Weed Science 39:474–479.

SchoLnbrunn E, Eschenburg S, Shuttleworth WA, Schloss JV, Amrhein N, Evans JN, Kabsch W (2001) Interaction of the herbicide glyphosate with its target enzyme 5- enolpyruvylshikimate 3-phosphate synthase in atomic detail. Proc Natl Acad Sci USA 98:1376–1380.

Schymkowitz J, Borg J, Stricher F, Nys R, Rousseau F, Serrano L (2005) The FoldX webserver: an online force field. Nucl. Acids. Res. 33:W382–W384.

Shen H, Jin D, Shu JP, Zhou XL, Lei M, Wei R, Shang H, Wei HJ, Zhang R, Liu L, Gu YF (2017). Large-scale phylogenomic analysis resolves a backbone phylogeny in ferns. GigaScience 7:p.gix116.

Soltani N, McNaughton K, Sikkema PH (2015) Field horsetail (*Equisetum arvense* L.) control in corn. Canadian Journal of Plant Science 95:983–986.

Taylor JA (1980) Bracken: an increasing problem and a threat to health. Outlook on Agriculture 10:298–304.

Ungaro A, Pech N, Martin JF, Mccairns RS, Mevy JP, Chappaz R, Gilles A (2017) Challenges and advances for transcriptome assembly in non-model species. PloS One 12:p.e0185020.

Vetter J (2009) A biological hazard of our age: Bracken fern [*Pteridium aquilinum* (L.) Kuhn]—A review. Acta Veterinaria Hungarica. 57:183–96.

Vilella AJ, Severin J, Ureta-Vidal A, Heng L, Durbin R, Birney E (2009) EnsemblCompara GeneTrees: Complete, duplication-aware phylogenetic trees in vertebrates. Genome Research 19:327–335.

Zeng L, Zhang Q, Sun R, Kong H, Zhang N, Ma H (2014) Resolution of deep angiosperm phylogeny using conserved nuclear genes and estimates of early divergence times. Nature communications 5:4956.

Zhang N, Zeng L, Shan H, Ma H (2012). Highly conserved low□copy nuclear genes as effective markers for phylogenetic analyses in angiosperms. New Phytologist 195:923–937.

Zhou Q, Liu W, Zhang Y, Liu KK (2007) Action mechanisms of acetolactate synthase- inhibiting herbicides. Pesticide Biochemistry and Physiology 89:89–96.

